# Machine Learning and Network Analysis to Predict Hypothetical Protein Functions of Aeromonas *hydrophila*

**DOI:** 10.1101/2025.07.22.666223

**Authors:** Harun Pirim, Zaidur Rahman, Saviz Saei, Sangam Gyawali, Mohammad Marufuzzaman, Nazanin Tajik, Hasan Tekedar

**Affiliations:** North Dakota State University, USA; University of Arkansas, USA; Mississippi State University, USA; Warner Bros. Discovery, USA

**Keywords:** Protein function prediction, Machine learning, Network analysis, A. hydrophilla

## Abstract

*Aeromonas hydrophila*, antibiotic resistant gram negative bacteria, is a major fish pathogen. Moreover, *A. hydrophila* is considered to cause 13% of gastroenteritis cases in the United States. Therefore, it is important to identify groups of proteins that are effective in antibiotic resistance and causing mortality in aquaculture. We train machine learning models on existing *A. hydrophila* genomes to predict functions of 83 carefully filtered hypothetical proteins. Network analysis is conducted to cluster these proteins based on their similarities. Both ML and network analysis inform about possible roles of these proteins in vaccine candidacy and fish mortality.

## 1. Introduction

The genus *Aeromonas* comprises a diverse group of facultatively anaerobic, gram-negative, motile bacilli or coccobacilli that naturally inhabit various aquatic environments. Aeromonads were initially classified as part of the *Vibrionaceae* family; however, advances in molecular phylogenetics, including the sequencing of 16S rRNA, 5S rRNA, and housekeeping genes, led to their reclassification into a distinct family, *Aeromonadaceae* Martinez-Murcia et al. (1992). Among the 36 identified species within the genus *Aeromonas*, 19 are considered emerging clinical pathogens Fernández-Bravo and Figueras (2020). However, over 95% of the reported clinical cases are associated with four significant species –*A. hydrophila, A. caviae, veronii*, and *A. dhakensis*. These species cause various human infections, from mild diarrhea and gastroenteritis to severe wound infections and fatal sepsis, particularly in immunocompromised individuals Truong et al. (2024); Greiner et al. (2021); Dharanendra et al. (2024); Takahashi et al. (2012); Bernabè et al. (2023). Aeromonads also infect a wide range of hosts, including amphibians, fish, reptiles, and mammals Kwon et al. (2019); Guo et al. (2024); Zhao et al. (2024). Notably, they are highly prevalent in farmed fish, where they cause diseases that result in high mortality rates and significant economic losses in the aquaculture industry Pei et al. (2021); Chen et al. (2019); Dubey et al. (2022).

Aquaculture, one of the fastest-growing sectors in global food production, provides a vital source of protein to meet the increasing demand for seafood for the growing world population. The U.S. aquaculture industry continues to expand, with an average sales per farm of $552,569 and total food fish sales reaching $819.6 million in 2023, marking a 14% increase from 2018 National Agricultural Statistics Service, USDA (2024). Catfish is a dominant species, accounting for 59% of all food fish sales in 2013, valued at $480.0 million. Sustainable aquaculture requires balancing profitability with responsible farming practices to mitigate environmental and disease-related challenges. However, increasing production demands have driven the adoption of intensive aquaculture systems that optimize water resource use. Intensive aquaculture practices, including high stocking densities, improper feeding rates, elevated temperatures, and poor water quality, create stressful conditions that weaken fish immune systems Wise et al. (2021). This increased stress heightens susceptibility to bacterial infections and facilitates disease spread. Opportunistic pathogens like *Aeromonas hydrophila* thrive in these environments, exploiting immunocompromised fish and causing severe outbreaks that compromise fish health and welfare, leading to significant economic losses for the aquaculture industry.

Hypervirulent *A. hydrophila* (vAh) is a primary pathogen of channel catfish and a causative agent of Motile Aeromonas septicemia (MAS) Joseph and Carnahan (1994); Pridgeon et al. (2014). MAS is a devastating disease characterized by hemorrhagic lesions, organ necrosis, and high mortality rates Tuttle et al. (2023); Zhao et al. (2019). These vAh strains can infect the fish despite lacking concurrent infections and were identified to be 200-fold more virulent than earlier *A. hydrophila* isolates Pridgeon et al. (2013). The disease can manifest rapidly from a few infected individuals to entire ponds within a few days, resulting in large-scale losses and the costs associated with disease management Zhang et al. (2020).

The pathogenicity of vAh is multifactorial and is attributed to key virulent elements and their genetic diversity, which limits the development of potential vaccines Zhao et al. (2020). Numerous virulence factors that make vAh highly pathogenic have been investigated. These include adhesins, cytotoxins, hemolysins, proteases, lipases, cell excitatory enterotoxin, pilins, flagellin, secretions systems, etc., which collectively enable it to adapt to diverse environments, evade host immune responses, and establish infections in numerous animals Dong et al. (2017); Allan and Stevenson (1981); Rasmussen-Ivey et al. (2016); Tekedar et al. (2024). Currently, antibiotic treatment remains the primary disease management strategy for *A. hydrophila* infections in catfish farming. Oxytetracycline is the only FDA-approved antibiotic for controlling *A. hydrophila* infections; however, emerging antibiotic-resistant *A. hydrophila* strains pose a growing concern Nhinh et al. (2021); Eid et al. (2022); Sherif and Kassab (2023); Yu et al. (2021); Adah et al. (2024). The heavy reliance on antibiotics has accelerated antimicrobial resistance (AMR), limited treatment efficacy, and raised concerns about the long-term sustainability of disease control strategies Cabello et al. (2016). While live attenuated vaccines have been successfully implemented against other bacterial pathogens in aquaculture, there is currently no licensed live attenuated vaccine for *A. hydrophila*.

Advances in genome sequencing have further illuminated the genetic landscape of *A. hydrophila*, revealing a multitude of hypothetical proteins—proteins predicted from genomic data but lacking functional annotation. Despite their enigmatic nature, these proteins are of profound interest, and exploring the function of these hypothetical proteins is essential for uncovering novel mechanisms of bacterial adaptation and virulence. Understanding these hypothetical proteins could also provide insights into combating antimicrobial resistance and developing effective strategies for managing *A. hydrophila* infections in aquaculture and clinical contexts. For example, recent studies have identified a hypothetical protein in *A. hydrophila* that facilitates colistin resistance, a concerning development given the critical importance of this antibiotic as a last-resort treatment for multidrug-resistant infections Liu et al. (2021).

Recent advances in bioinformatics have introduced various tools that facilitate the functional annotation of hypothetical proteins. These tools leverage domain, family, and ontology databases to predict protein functions, enhancing our understanding of bacterial biology. Utilizing these tools in conjunction with machine learning algorithms, this research aims to predict and analyze the functions of hypothetical proteins in *A. hydrophila*. Table 1 lists some of the bioinformatics tools employed in this study.

**Table 1:** Bioinformatics tools to characterize proteins.

| Bioinformatics Tool | Description |
| --- | --- |
| CATH | A classification of protein domains based on sequence information, structural, and functional properties (Das et al., 2015) |
| Pfam | A complete and accurate classification of protein families and domains (Finn et al., 2007) |
| DEG | Identifies essential genes likely to be common to all cells (Zhang et al., 2004) |
| PSORTdb | Determines the sub-cellular localization of the protein (Nakai and Horton, 1999) |
| BLAST | A similarity searching program for homogenous DNA sequences and proteins based on the excess sequence closely matched (Altschul et al., 1990) |

One of the latest trends in proteomics is the advent of machine learning (ML) and deep learning (DL) particularly. Proteomics data is high-dimensional, making it well-suited for these computational methods. Studies on ML for protein function prediction (Barla et al., 2008; Swan et al., 2013; Bouwmeester et al., 2020; Neely et al., 2023; Vishnoi et al., 2020) suggest that we can enhance protein related prediction tasks. Although DL offers combining complex features and infusing large datasetsArmah-Sekum et al. (2024); Boadu and Cheng (2024), it requires more computational resources and assumes abundance of data availablity.

(Bonetta and Valentino, 2020) review protein function prediction algorithms and feature selection methods that consider biomedical text-derived features in addition to physicochemical and amino acid composition features. They report the success of traditional ML algorithms, that makes them a viable tool, compared to deep learning approaches. One particular reason of this success can be related to the features selection and concatenation that produce a tabular dataset for ML. (Yu and Luo, 2023) develop a framework to predict peptide and protein functions using ensemble algorithm such as random forest, XGBoost and AdaBoost. They report the framework’s superior performance compared to the state of the art methods. We apply the same ensemble algorithms to predict multilabel GOterms. Multilabel classification allows a protein sequence to be assigned many GO-terms instead of a single one.

By utilizing these tools and machine learning algorithms, this research aims to predict and analyze the functions of hypothetical proteins found in *Aeromonas hydrophila*. The insights gained will not only enhance our understanding of the pathogenic mechanisms of these bacteria but also potentially lead to the development of more effective treatment strategies. Ultimately, this work supports the sustainability and productivity of the catfish farming industry by providing new avenues to combat bacterial infections.

## 2. Methodology

The methodology starts with data collection and feature selection followed by machine learning for predicting protein function.

### 2.1. Data Collection and Feature Selection

To predict the functions of hypothetical proteins in Aeromonas bacteria, we begin by assembling a comprehensive dataset of proteins with known functions and their associated features. We then employ three machine learning algorithms—Random Forest, XGBoost and Adaboost to train on this dataset. The model learns to recognize complex patterns and correlations between the features and the known protein functions. Once the models are trained and validated, they can predict the functions of new, uncharacterized hypothetical proteins by analyzing their features within the learned framework.

#### 2.1.1. Protein Sequence Data and GO terms

Under the *Aeromonas* genus, protein sequences of other species including *hydrophila* are used for the proposed prediction model. According to the last update till 2023, Uniprotkb (Consortium, 2018) has exactly 427837 annotated *Aeromonas* sequences including both reviewed and unreviewed sequences. Uniprotkb uses annotation score as the level of annotation and the quality of the information for each protein entry in the database. The annotation score ranges from 1 to 5 stars. The higher the score, the more information and higher quality annotation the protein entry has.

For this study, protein sequences with annotation score 3, 4 and 5 was selected to maintain a balance between the number of protein sequence and the quality of the annotation. This totaled to 31321 protein sequences.

The aim of the suggested model is to predict the functions linked to protein sequences, which are expressed as Gene Ontology (GO) terms. Gene Ontology is a standardized system used to describe the roles of genes and their associated proteins in a consistent way across species. Protein functionality is categorized into several interconnected levels, aligning with the three primary GO categories: molecular function, biological process and cellular component (Lee et al., 2007). Molecular refers to the basic activities of proteins at a molecular level, such as binding or catalysis. Biological Process describes larger processes in which proteins are involved, like signaling pathways or cell growth. And the Cellular Component describes the location within a cell where the protein typically operates, such as the nucleus, membrane, or mitochondria.

Because proteins can have complex and varied roles, a single protein sequence can be associated with one or more GO terms, reflecting its different functions across the molecular, biological, or cellular levels. This makes the problem of predicting protein function a supervised multi-label classification task. In multi-label classification, unlike single-label classification where each instance belongs to just one category, each protein can belong to multiple GO categories simultaneously.

The GO system is organized hierarchically, with broad, general terms (parent terms) at the top, and more detailed, specific terms (child terms) at the bottom. In this work, the dataset has been curated by selecting the child GO terms. Child GO terms provide more precise information about the function of a protein. While parent terms offer general descriptions, child terms capture the specific biochemical activities, biological processes, or cellular components involved. In biological research, it is often more valuable to know exactly what a protein does at the most granular level. Selecting child GO terms enables the model to offer these insights, rather than vague, high-level descriptions that could apply to many proteins. For example, knowing that a protein is involved in a specific kinase pathway is more useful than simply knowing it has general enzyme activity.

One of the main drawbacks of selecting child GO Terms is that focusing on child terms significantly increases the number of target columns in the dataset, as each child term represents a distinct, granular function. This expanded set of target labels adds complexity to the model, making it harder to train and potentially increasing the risk of overfitting, especially when dealing with limited data. The increased complexity could also lead to higher computational costs, both in terms of time and resources, when processing large datasets with numerous target labels.

The GO terms used for this study were collected directly from the UniProtKB database. Among the 31321 sequences, 1540 sequences’ entries were deleted from the UniprotKB database and had no go terms. Those sequences were removed which resulting the total sequence number of 29,781.

#### 2.1.2. Features

The features of the main dataset are divided into four main categories.

##### Database of Essential Genes (DEG)

This database, introduced by Zhang et al. (2004), catalogs genes essential for the survival of various organisms. It’s divided into three sections: Bacteria, Archaea, and Eukaryotes, each representing a unique life domain based on cellular and genetic characteristics. Users can perform BLAST searches, comparing query sequences with essential gene sequences in DEG. Initially, protein BLAST is conducted against each DEG section, and two types of features are extracted from these search results MaxID, representing the maximum percent identity among the hits and TotalHits the total number of hits in the database.

These features were determined for each of the three DEG databases, giving a set of six features for each protein sequence.

- DEGBacteria_MaxID: Maximum percent identity for Bacteria Essential Gene Database
- DEGBacteria_TotalHits: Total number of hits Bacteria Essential Gene Database
- DEGArchea_MaxID: Maximum percent identity for Archaea Essential Gene Database
- DEGArchea_TotalHits: Total number of hits Archaea Essential Gene Database
- DEGEukar_MaxID: Maximum percent identity for Eukaryotes Essential Gene Database
- DEGEukar_TotalHits: Total number of hits Eukaryotes Essential Gene Database

Sequence similarity, indicated by the MaxID attribute, often suggests functional similarity in biological sequences. For example, a protein closely matching a gene in the DEG database might share a similar essential role. MaxID values across life domains can hint at a protein’s function conservation: high MaxID in all domains suggests a universal function, while high MaxID in one domain points to a domain-specific role. The TotalHits feature indicates the protein family size, with larger families implying broader functions and smaller families suggesting specialized roles.

The DEG database only supports single protein sequence searches at a time. Querying all 29,781 sequences in DEG would require 89,343 searches, making manual querying impractical. Lacking an API for bulk queries, a Python web scraping script was developed for automating this process, as described in figure 1.

**Figure 1:**
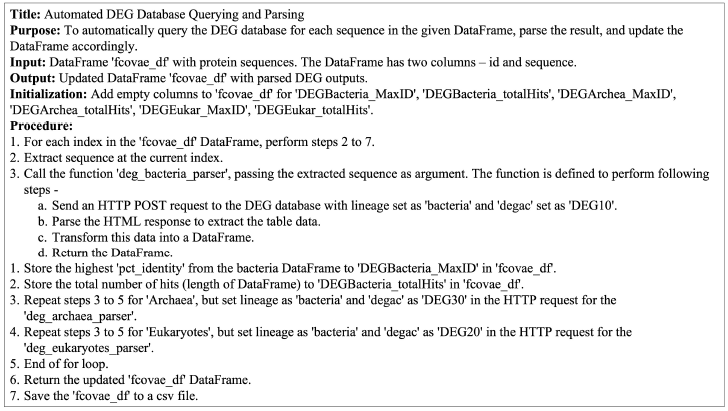
DEG Database Query Algorithm

The finding that most matched essential genes are from the Bacteria group, among Bacteria, Archaea, and Eukaryotes, makes sense since the sequences were taken from a bacterium. This result highlights the genetic similarities within a group and the distinct differences between life’s domains.

##### PSORTb Localization

The protein sequences are then analyzed using PSORTb v3.0.2, the most precise bacterial localization prediction tool available as of 2010 (Yu et al., 2010). PSORTb predicts where in a bacterial cell a given protein is likely to be located. From the PSORTb analysis, six features were collected for each protein sequence.

- Cytoplasmic (Score): Indicates probability that the protein is located in the cytoplasm of the cell.
- CytoplasmicMembrane (Score): Represents the likelihood of the protein being in the cytoplasmic membrane.
- Periplasmic (Score): It is the probability of the protein being in the periplasm, the space between the outer membrane and the cytoplasmic membrane in Gram-negative bacteria.
- OuterMembrane (Score): This is the probability that the protein is located in the outer membrane.
- Extracellular (Score): This score indicates the likelihood of the protein being located outside the cell. Although, for a single-celled bacterium, it is not possible for any protein to be located outside of the cell.
- Final Prediction: This is the subcellular location that PSORTb predicts with the highest probability. This will serve as a categorical feature in the final dataset. Final Prediction has any one of the above 5 categories or an “unknown” category.

PSORTb team has a standalone version of the PSORTb in a Docker image for local installation with pre-built dependencies. The docker image were run in a local machine to parse the localization predictions for all sequences. PSORTb was able to return localization information for all the 29,781 sequences.

##### NCBI BLAST

The National Center for Biotechnology Information (NCBI) is a part of the United States National Library of Medicine (NLM), a branch of the National Institutes of Health (NIH). The NCBI maintains a unique protein sequence repository known as the non-redundant (nr) (Pruitt et al., 2005) database. This database amalgamates non-redundant sequences from GenBank CDS translations and various specialized sources. Its “non-redundant” designation stems from its method of merging identical sequences into a single entry. Utilizing BLASTP (Mahram and Herbordt, 2015), a protein comparison tool, researchers can align query sequences against this expansive collection of protein sequences from diverse organisms, offering insights into the sequence’s evolutionary relationships with other proteins.

For comprehensive analysis, each sequence underwent a BLASTP search. To balance scope and data manageability, the maximum hits per sequence were capped at 300. These 300 matches were first filtered based on query coverage, which assesses the extent to which the query sequence aligns with database sequences. Priority was given to hits enveloping the entire query sequence, initially selecting those with 100% coverage. In cases where no hits achieved complete coverage, those with the highest available coverage were chosen.

The hits were filtered further by the percent identity. It depicts similarity level of the aligned sequences. All the hits with 80% or more identity were selected for this study. If any sequence did not have any hits with 80% or higher identity percentage, the hits with maximum percentage were selected. Upon filtering, two features were kept for each sequences.

- count_blast: The number of hits that passed the filtering steps for each sequence. This can suggest functional conservation across these sequences, as sequences with similar structure often perform similar functions.
- AvgIdBlast: The average percent identity of the filtered hits for each sequence. The higher the identity, the more likely the proteins share similar functions, as evolutionary pressure tends to conserve functionally important sequences.

NCBI has a command-line tools to run BLAST called BLAST+ (Camacho et al., 2009). As the whole search operation requires a lot of computational resources and time, it was performed on the High-Performance Computing cluster, managed by Center for Computationally Assisted Science and Technology (CCAST) at North Dakota State University. A local version of nr database was downloaded and BLAST+ was executed with an input of the protein sequence fasta file, which performed the queries against the downloaded local nr database. Out of 29,781, 1 sequence did not have any blast hits, resulting in total 29,780 sequences.

##### Physicochemical Features

The set of features comprises two main components: the calculation of a protein’s biophysical characteristics using the “ProteinAnalysis” class within BioPython, and the determination of various sequence-based attributes utilizing the PyPro library. The ensuing enumeration details the features specific to both BioPython and PyPro.

- Molecular Weight: It is the sum of the atomic weights of all the amino acids and any other atoms in the protein.
- Instability Index: This is a measure of the protein’s stability in a test tube environment. Proteins with an index less than 40 are predicted to be stable, and those with an index greater than 40 are predicted to be unstable.
- Isoelectric Point: The pH at which a particular molecule or surface carries no net electrical charge. This is important for protein function as it affects protein structure and interactions.
- GRAVY (Grand Average of Hydropathy): This is the average hydropathy index of all the amino acids in a protein. A higher GRAVY index usually indicates that the protein is hydrophobic and likely to be found within cell membranes.
- Secondary Structure Fraction: These are the fractions of amino acids which tend to be in helices, turns, or sheets in the secondary structure of the protein. Proteins fold into these secondary structures. This feature has three values, one for each type of secondary structure (helix, turn, and sheet).
- Extinction Coefficient: This represents how much light a protein absorbs at a certain wavelength. It’s commonly used to estimate protein concentration. Two values are returned: one assuming all pairs of Cys residues form cystines, and one assuming all Cys residues are reduced.
- Grouped Amino Acid Composition: This represents the occurrence frequencies of each type of amino acid. It has 20 values, one for each amino acid.
- CTD (Composition, Transition, and Distribution) descriptors: These are used to characterize the composition, transition, and distribution of amino acids within a protein sequence. It has 147 values.

The physicochemical characteristics and amino acid composition of proteins play a crucial role in shaping their threedimensional structure, which is essential for their functional capabilities. Knowledge of these attributes enables predictions about a protein’s behavior and interactions within a biological context. Additionally, examining grouped amino acid composition and CTD descriptors offers detailed insights into the protein’s sequence. Such features are instrumental in identifying specific patterns or motifs that are intimately connected to the protein’s function.

Among all the sequences, 5 had “X” in them, which represents unknown amino acid. For those sequences, it was not possible to get the accurate physicochemical features. Those sequences were removed and the number of total sequences updated to 29,775.

### 2.2. Machine Learning for Predicting Protein Function

The rapid growth of biomedical research and biotechnology has led to an explosion of biomedical data (Cremin et al., 2022). To extract useful knowledge from this data, researchers are increasingly turning to machine learning. In the field of bioinformatics, machine learning algorithms are used to discover predictable patterns in biological systems, thus helping to predict the function of hypothetical proteins (Yang et al., 2020; Camacho et al., 2018). Several machine learning techniques are used for this purpose, as summarized in Table 2.

**Table 2:** ML Algorithms used in bioinformatics.

| Machine Learning Techniques | Description | Limitation |
| --- | --- | --- |
| Random Forest | An ensemble classification model that uses a majority vote among trees for prediction (Breiman, 2001) | The algorithm is slow and ineffective for a large number of trees (Oshiro et al., 2012) |
| Gaussian Naive Bayes | A classification model that assumes a simple Gaussian distribution for data from each label (Zhang, 2005) | The model becomes infeasible with increasing features due to the independence assumption (Wickramasinghe and Kaluturage, 2021). |
| AdaBoost | A robust classification method that combines many weak learning classifiers to create a strong one (Bartlett et al., 1998) | Sensitive to outliers and noisy data (Tanha et al., 2020). |
| XGboost | An ensemble classification with multiple weak classifications in parallel tree boosting (Chen and Guestrin, 2016) | Sensitive to outliers (Chao et al., 2023). |
| MLP (Multi-Layer Perceptron) | A feedforward artificial neural network with a non-convex loss function (Tolstikhin et al., 2021) | Different random weight initializations lead to different validation accuracy (Glorot and Bengio, 2010). |

K-fold cross-validation, a critical method in machine learning, is used to balance bias and variance in performance estimation, avoiding skewed results towards any data subset. It involves dividing the data into ‘k’ sets, using each as a test set while the others serve as training data, and then averaging the model’s performance across these sets. This approach, particularly the 10-fold cross-validation, is a popular choice for its robustness, as seen in various studies (Martin et al., 2005; Dor and Zhou, 2007). In this study, a 10-fold cross-validation is adopted to evaluate three machine learning models.

Six evaluation measures – Accuracy, Precision, Recall, F1Score, AUC-ROC and Log Loss are collected for each trained model. For all the three models, these scores represent the average values for all the train-test splits in 10-Fold CrossValidation. Following sections discuss these scores in detail.

#### 2.2.1. Accuracy

Accuracy in classification tasks quantifies the ratio of correct predictions to total predictions made by a model. Typically, in binary or multiclass classification, accuracy is straightforward, calculated as the number of correct predictions divided by the total number of predictions.

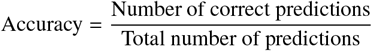

However, in multi-label classification, where an instance may simultaneously belong to several categories, accuracy takes a different form. Here, it’s measured as subset accuracy, also known as “exact match”. This metric considers a prediction accurate only if every label of an instance is correctly identified. Essentially, the set of predicted labels must precisely align with the actual set of labels for a perfect accuracy score.

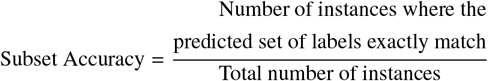

In practical terms, accuracy is often evaluated using a test dataset, which is a randomly selected portion (like 20%) of the main dataset. This helps in understanding how the model performs on data it hasn’t seen before.

#### 2.2.2. Precision

Precision in machine learning, especially within classification tasks, serves as a key metric of a model’s accuracy in predicting positive outcomes. Essentially, it’s the ratio of true positives (TP) to the sum of true positives and false positives (FP). This ratio indicates how often the model’s positive predictions are actually correct. A high precision score suggests that a model’s positive predictions are predominantly accurate.

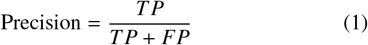

In the context of multi-label classification, where instances may belong to multiple categories, calculating precision is a bit more complex. A commonly used approach is the sample-average precision. This technique calculates precision for each instance separately, then averages these scores across all instances. This approach offers a nuanced view of the model’s precision, especially in scenarios involving multiple labels. The precision scores for different models are typically compiled in table 4 for comparative analysis, particularly using test data to assess model performance..

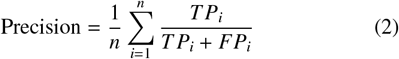

Where, *n* = number of instances (samples) in the test dataset

#### 2.2.3. Recall

The recall score, often referred to as sensitivity or the true positive rate, plays a crucial role in evaluating the effectiveness of a classification model. This metric primarily focuses on the model’s ability to correctly identify actual positive instances. In essence, recall addresses the question, “Of all the actual positives, how many has the model successfully detected?” The calculation for recall involves dividing the number of true positives (TP) by the total of true positives and false negatives (FN), which represents instances that are positive but were missed by the model. The formula can be expressed as:

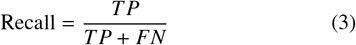

In situations involving multi-label classification, where an instance can belong to multiple categories, the calculation of recall requires a bit more nuance. It involves determining the recall for each instance individually and then calculating the average of these recall scores across all instances. This approach provides a more detailed understanding of how well the model performs in correctly identifying positive instances across a range of different labels.

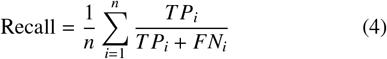

Where, n = Number of instances (samples) in the test dataset

#### 2.2.4. F1-Score

The F1 score is a balanced metric used to evaluate the accuracy of a model, blending aspects of both precision and recall. Precision measures the proportion of actual positives among the model’s positive predictions, while recall assesses the proportion of actual positives correctly identified by the model. The F1 score harmonizes these two elements, offering a more comprehensive view of the model’s performance than either metric alone. It’s calculated as the harmonic mean of precision and recall, achieving its optimum at 1 (perfect precision and recall) and its minimum at 0. In scenarios involving multi-label classification, the F1 score is determined for each label separately, and then these scores are averaged to yield an overall measure. This methodology offers a nuanced understanding of the model’s effectiveness across different categories.

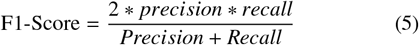

#### 2.2.5. AUC-ROC and Log Loss

The AUC-ROC curve is a critical metric for evaluating the effectiveness of a classification model across different threshold levels. The ROC (Receiver Operating Characteristics) curve visualizes the trade-off between the true positive rate (TPR) and the false positive rate (FPR), offering insights into how well the model distinguishes between the two classes under various thresholds. The AUC (Area Under the Curve) quantifies this ability, representing how effectively the model can differentiate between the classes. A higher AUC value indicates a model’s superior capability in correctly classifying the negatives (0s) as negatives and the positives (1s) as positives, essentially reflecting its discriminative power. The AUC-ROC curve is generated by plotting equation 6 on x axis and equation 7 on y axis.

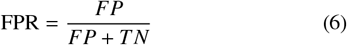

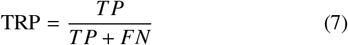

#### *TP* + *FN*

Where, TP = True Positives, FP = False Positives, FN = False Negatives and TN=True Negatives

In multi-label classification, AUC-ROC is calculated for each label and the average of the scores over the labels are the final AUC-ROC score. On the other hand, Log Loss (Logarithmic Loss) is a loss function that is used in machine learning to evaluate the predictions of probabilities of membership to a given class. The equation 8 is for Log Loss in binary classification.

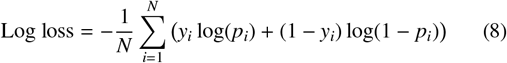

Where,

- *N* is the number of observations,
- Σ is the sum over all observations,
- *y* is the true class (0 or 1),
- *p* is the predicted probability that the observation is of class 1.

The smaller the Log Loss, the closer the predicted probabilities are to the true classes, which indicates better model performance. In multi-label classification, the log loss calculation method is similar to AUC-ROC method. The log loss is calculated for each label, and then the results are averaged. This treats each label as an independent binary classification problem.

#### 2.2.6. Vaccine Development Candidacy using GO Terms

The identification of vaccine candidates is a crucial step in the development of effective vaccines. And the identification fundamentally relies on understanding the biological functions of proteins within a pathogen. Predicting protein functions allows us to target proteins that are essential for the pathogen’s survival, virulence, or interaction with the host immune system. Proteins involved in critical biological processes, are more accessible to the immune system and are prime targets for vaccine development. By associating proteins with specific GO terms, we can infer their roles in pathogenicity and their potential to generate an immune response. This approach streamlines the discovery of vaccine candidates by narrowing down the list to proteins most likely to be effective, saving time and resources in experimental validation. By leveraging predicted functions through GO terms, we aim to systematically identify proteins most suitable for vaccine development, maximizing the utility of computational predictions in guiding experimental efforts.

For each hypothetical protein, the machine learning models provided sets of predicted GO terms encompassing molecular functions, biological processes, and cellular components. For each prediction method, we collected the unique GO terms associated with the proteins predicted by that method. This resulted in separate sets of GO terms corresponding to each prediction approach. Each set of unique GO terms was individually filtered using a predefined list of keywords pertinent to vaccine development, such as “outer membrane”, “adhesion”, “surface antigen”, “toxin”, “virulence”, “pathogenicity”, “invasion”, “secretion system”, “immune evasion”, “antigen”, “epitope”, “immune response”, “innate immunity”, “membranebound”, “type III secretion system”, “type IV secretion system”, “activation of immune response”, “positive regulation of defense response”, “response to bacterium”, “defense response to virus” and “innate immune response”. This filtering process aimed to isolate GO terms that indicate a protein’s potential to elicit an immune response or play a role in pathogenic mechanisms. After filtering, the selected GO terms from each prediction method were mapped back to the corresponding hypothetical proteins. This mapping allowed us to identify specific proteins associated with functions relevant to vaccine development within each prediction set. Proteins linked to the filtered GO terms were considered potential vaccine candidates due to their predicted involvement in immunologically significant processes.

## 3. Results and Discussion

The effectiveness of different models can be compared using their accuracy scores (Table 3).

**Table 3:** The accuracy scores of three trained models Table 5: The recall scores of the models.

| Models | Accuracy |
| --- | --- |
| Random Forest | 87.44% |
| XGBoost | 86.63% |
| AdaBoost | 78.65% |

**Table 4:** The precision scores of the models.

| Models | Precision |
| --- | --- |
| Random Forest | 96.55% |
| XGBoost | 97.29% |
| AdaBoost | 96.60% |

**Table 5:** The recall scores of the models.

| Models | Recall |
| --- | --- |
| Random Forest | 93.97% |
| XGBoost | 94.78% |
| AdaBoost | 93.30% |

The Random Forest model demonstrates robust predictive capabilities, correctly identifying the entire label set for nearly 92.84% of the instances. The XGBoost model edges out slightly ahead, with its accuracy reaching 93.58%, indicating its slightly superior efficiency in accurately predicting labels. On the other hand, the AdaBoost model shows a comparatively lower accuracy of 73.38%, suggesting it’s less effective in predicting the correct labels across instances compared to the other two models.

The Random Forest model exhibits a slight edge over XGBoost, achieving a precision of 99.04%. This subtly higher score suggests that Random Forest is marginally more accurate in correctly identifying positive cases, indicating its slight superiority in precision compared to XGBoost, which is close behind with a precision of 99.02%. Both models showcase exceptional precision, implying that their predictions are almost invariably correct.

AdaBoost, while demonstrating a lower precision score of 95.77%, still maintains a commendable level of accuracy. This score, though not as high as the other two models, indicates that AdaBoost, despite a higher tendency for false positives, generally performs well in accurately identifying positive instances. However, it’s clear that in terms of precision, Random Forest and XGBoost lead the way, with their near-flawless performance in this metric.

Reflecting on the recall scores from the table, the performance of the three models shows noticeable differences. The Random Forest model demonstrates a solid recall rate of 96.98%, indicating its proficiency in correctly identifying a high percentage of positive instances. This suggests that the model is quite adept at capturing relevant data, missing only a small fraction of positive cases. XGBoost, with a recall score of 97.74%, surpasses Random Forest, showcasing a slightly higher effectiveness in identifying positive instances. This score reflects its superior capability in recognizing relevant data, making it a highly reliable model in terms of recall.

In comparison, AdaBoost lags somewhat behind with a recall of 93.85%. Although this is a respectable score, it indicates that AdaBoost is somewhat less efficient in capturing all relevant positive instances compared to the other two models. While it successfully identifies a majority of the positives, its performance in this aspect is not as high as that of Random Forest or XGBoost. This table clearly delineates the recall capabilities of each model, highlighting their respective strengths in identifying positive instances.

Table 6 listed the F1-scores on test dataset for all three models.

**Table 6:**
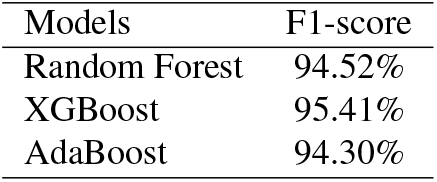
F1-score values of the models.

| Models | F1-score |
| --- | --- |
| Random Forest | 94.52% |
| XGBoost | 95.41% |
| AdaBoost | 94.30% |

In a comparison of three models using the F1-score, XG-Boost excelled with a leading score of 98.13%, demonstrating its efficient balance of precision and recall. This performance is likely due to its gradient boosting framework, which effectively handles diverse data types and imbalances. The Random Forest model also performed well, achieving an F1-score of 97.57%. Known for its resistance to overfitting and aptitude for highdimensional data, this model’s slightly lower score compared to XGBoost still reflects its robust predictive capability. AdaBoost, with an F1-score of 93.26%, ranked lowest among the three. While it’s a strong boosting technique, in this test, it didn’t match the performance of the other two models, indicating potential limitations in precision or recall. This comparison highlights the importance of choosing the right model based on dataset characteristics and the specific problem.

Both AUC-ROC and Log Loss scores are calculated for the on the test dataset for all three models. The scores are listed in Table 7.

**Table 7:** AUC-ROC and Log Loss scores of the models.

| Models | AUC-ROC | Log Loss |
| --- | --- | --- |
| Random Forest | 0.9978 | 0.00195 |
| XGBoost | 0.9982 | 0.00141 |
| AdaBoost | 0.9981 | 0.06624 |

### 3.1. Prediction on Hypothetical proteins

The hypothetical protein dataset has 83 protein sequences without any target GO terms. Those sequences were fully unrecognized by PANNZER (Koskinen, TÖrÖnen et al. (2015) model. The GO terms of those sequences were predicted by the three models trained with Random Forest, XGBoost and AdaBoost. According to the proposed hypothesis, it was expected to get the GO term prediction of at least 10% of the hypothetical proteins (2015 proteins). But surprisingly, All three models could get the GO terms predictions for all the hypothetical proteins. Table 8 summarizes the result.

**Table 8:** Hypothetical protein prediction summary.

| Models | Number of Predictions | Prediction Percentage |
| --- | --- | --- |
| Random Forest | 83 | 100% |
| XGBoost | 83 | 100% |
| AdaBoost | 83 | 100% |
| PANNZER | 65 | 78.31% |

The frequency of the predicted GO Terms for the hypothetical proteins by all these three models are presented in the figure 2, 3 and 4.

**Figure 2:** Frequency of the GO terms in the final dataset

**Figure 3:**
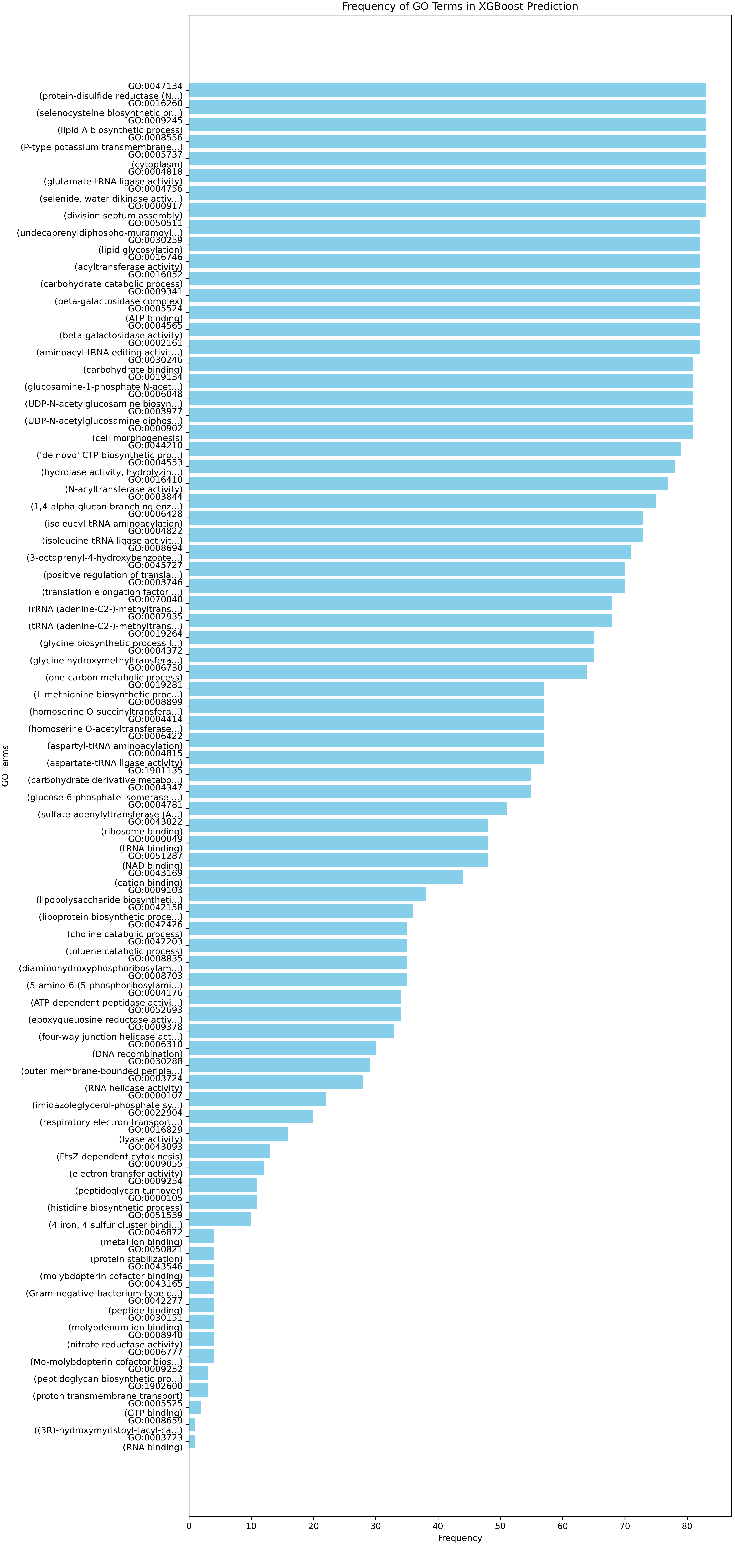
Frequency of the GO terms in the final dataset

**Figure 4:**
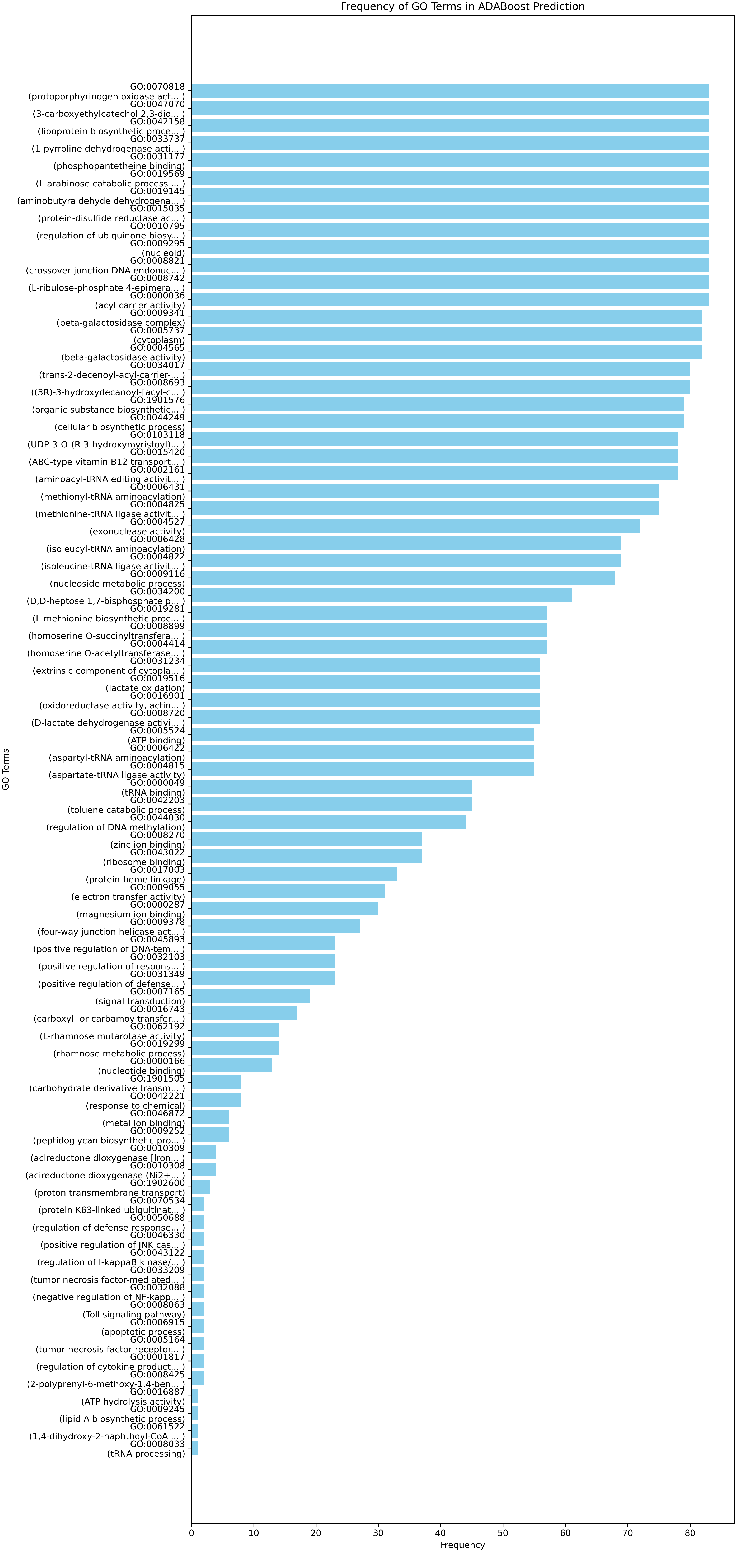
Frequency of the GO terms in the final dataset

The number of unique GO terms predicted by XGBoost algorithm is higher with 21 GO terms. The Random Forest and Adaboost predictions are lower with 14 and 11 unique GO terms. Although the frequency distributions are different in all three algorithms, two highly frequent GO terms are common in all three predictions GO:0005575 (Cellular Component) and GO:0006629 (Lipid Metabolic Process).

### 3.2. Feature Importances

Feature importance refers to a technique used in predictive modeling to assign a score to input features based on how useful they are at predicting a target variable (Zien et al., 2009). It measures the relative contribution of each feature towards the model’s performance, allowing us to understand which features have the most influence on the predictions made by the model. This understanding is crucial because, it helps in model interpretation, helping to uncover the underlying patterns that the model is leveraging for predictions.

#### 3.2.1. Random Forest

Each decision tree within the Random Forest is constructed through a process that involves selecting random subsets of features and instances. This randomness introduces diversity among the trees, contributing to the robustness of the model. During the construction of these trees, the improvement in the model’s performance, attributable to each feature, is quantified at each node where a split occurs.

The feature_importances_ attribute of the Random Forest model encapsulates the essence of this methodology by aggregating the individual contributions of each feature across all the trees in the forest. Specifically, it computes the average decrease in the chosen criterion that results from splits over a particular feature. Consequently, features that frequently contribute to homogenizing the target variable within the subsets created by splits are deemed more important.

To calculate the overall feature importances in a Random Forest model, the algorithm follows three main steps. Firstly, For each decision tree, it determines the total decrease in the splitting criterion attributable to each feature. This is done by summing the decreases across all nodes in the tree where the feature is used to split the data. Secondly, it normalizes these totals by the number of decision trees in the forest to obtain the average importance of each feature. Thirdly, it ranks the features based on their average importance to identify which features contribute most to the prediction accuracy of the model. Figure 5 shows the top 35 features, which have greater than 0.007 importance score.

**Figure 5:**
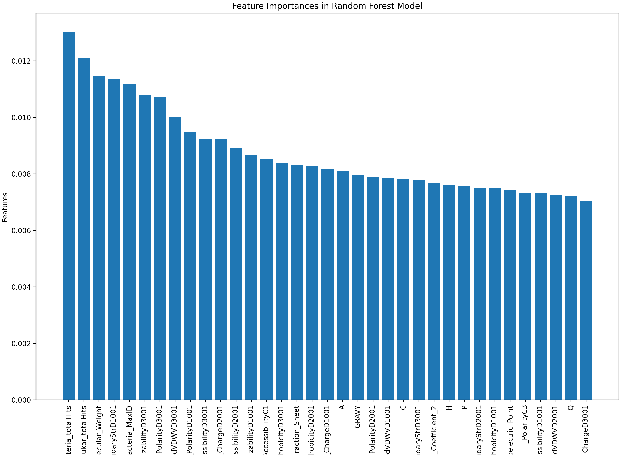
Random Forest Aggregated Feature Importances (Importance > 0.007)

#### 3.2.2. XGBoost

The XGBoost model comprised multiple estimators, each contributing to the final decision-making process. For each estimator within the ensemble model, feature importances were extracted using the feature importances attribute provided by the sklearn library. This attribute quantifies the contribution of each feature to the model by computing the relative importance, which is measured based on how much each feature splits and reduces the impurity across all trees in the model. This method aggregates these values across all estimators to ensure a comprehensive evaluation of each feature’s impact.

The accumulated importances were then averaged over the number of estimators in the model, yielding an average importance score for each feature. This averaging ensured that the derived importance metrics were representative of the model as a whole, rather than being biased towards any single estimator. After calculating the average importance scores, these scores were associated with their corresponding feature names. All the features that have more than 0.007 importance were plotted in figure 6. Among all the features, 44 features have high importance scores.

**Figure 6:**
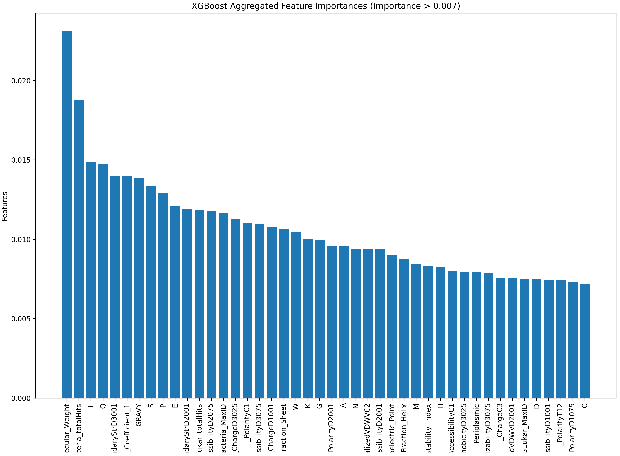
XGBoost Aggregated Feature Importances (Importance > 0.007)

#### 3.2.3. AdaBoost

The AdaBoost algorithm utilizes a series of estimators, each playing a role in the ensemble’s overall decision-making process. These importances are determined using the similar feature importances attribute in the AdaBoost estimators, which calculates how each feature reduces the weighted impurity in decision trees, thereby reflecting each feature’s ability to improve model accuracy.

Importance scores were accumulated for each feature across all estimators and subsequently averaged to provide a representative measure of importance across the ensemble. This method of averaging helps mitigate biases that might arise from any single estimator’s dominant influence. The averaged importance scores were then mapped to their corresponding feature names. Features were sorted based on their importance scores in descending order, and the top features were visualized through the figure 7. According to the result, 36 top features has more than 0.007 importance score.

**Figure 7:**
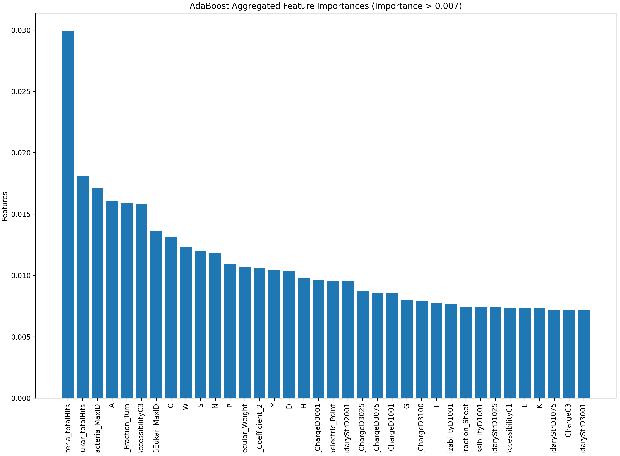
AdaBoost Aggregated Feature Importances (Importance > 0.007)

All the feature importance scores for all three algorithms were combined in figure 9. From this figure, importance score for different groups can easily be compared for the three algorithms. It can be seen from figure 9, NCBI features have the lowest importance score for all three algorithms. And DEG features have comparatively higher importance scores. Physicochemical features have higher importance in Random Forest and XGBoost algorithm. But those have lower importance scores in AdaBoost algorithm.

**Figure 8:**
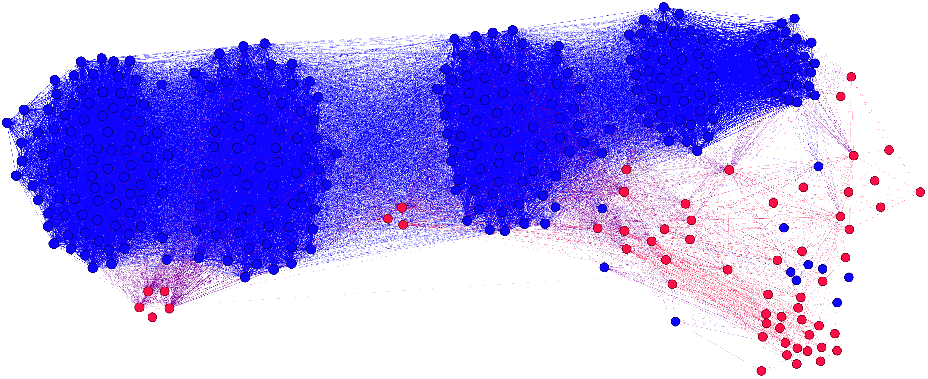
The similarity network of proteins. Red nodes represent hypothetical proteins.

**Figure 9:**
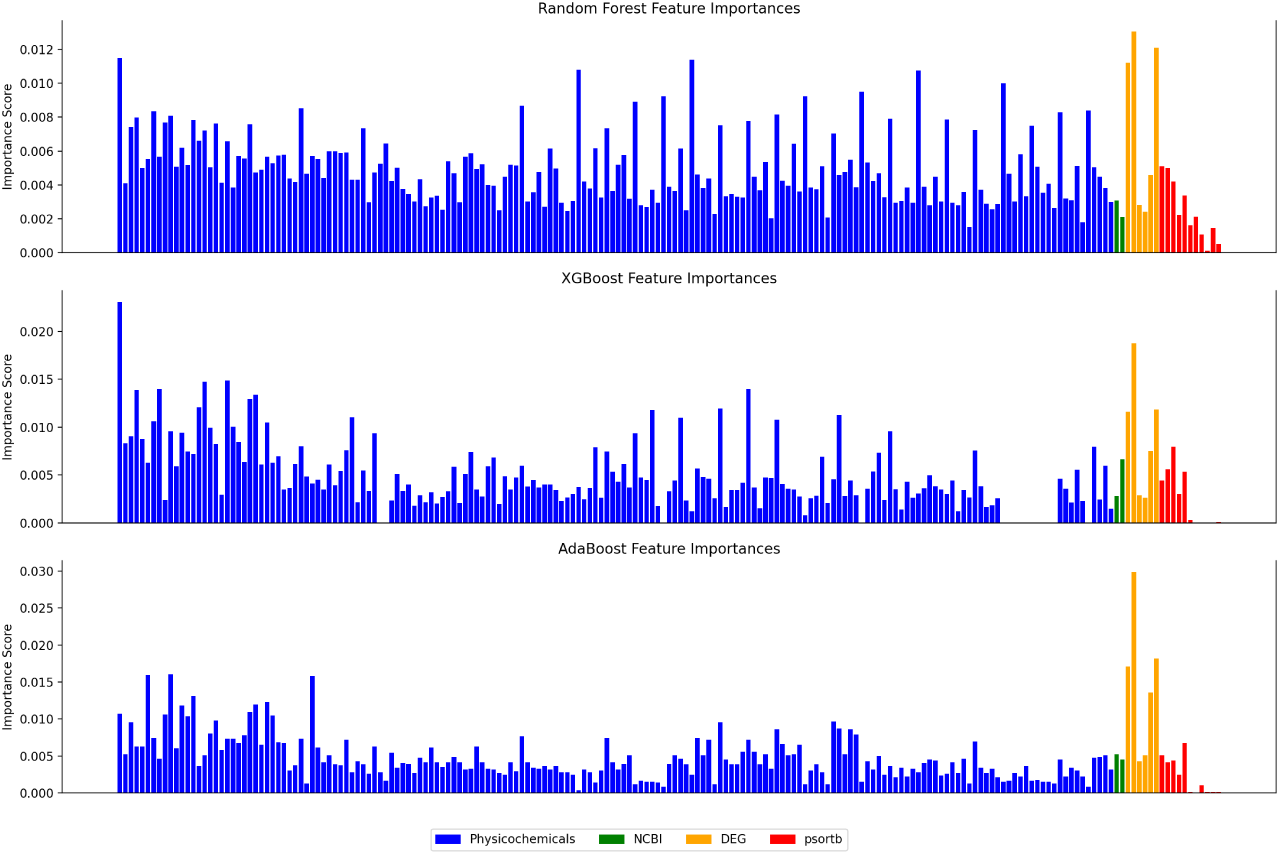
Feature importance scores for all three algorithms

### 3.3. Network Analysis for Prediction

In addition to using ML to predict hypothetical protein functions, we constructred a similarity network of proteins that involves both hypothetical and regular ones. We filtered our regular proteins based on their GO term descriptions. We kept the ones that has at least two distinct GO terms that includes one or more of these keywords: “immune”, “antigen”, “pathogen”, “cytokine”, “receptor”, “binding”, “membrane”, “extracellular”. These keywords represent vaccine candidacy of proteins based on their relevance to immune response, acting as an antigen, and pathogen interaction. The filter reduced our regular proteins of focus from 29,775 to 385. We merged them with 83 hypothetical proteins to construct the similarity network using 195 features. Gower distance matrix (GDM) is calculated followed by conversion to the similarity matrix using [1 − *GDM*] where 1 is the matrix of the same size with GDM filled with ones. In order to clarify network relationships we set the threshold of 75th percentile of the similarity scores, i.e. 0.875, trimming the edges lower than this score. The resultant network has 468 nodes with 27,144 edges. We focused on the largest connected component of the network that has 366 nodes including 59 hypothetical proteins shown in Figure 8.

To predict GO terms for these 59 hypothetical proteins, we examined their connections to regular proteins within the largest connected component of the network. Specifically, for each hypothetical protein, we identified its connected regular proteins and extracted the GO terms associated with those proteins. Out of the 59 hypothetical proteins, 12 did not have any connections to regular proteins. For the remaining 47 hypothetical proteins, we treated the GO terms of their connected regular proteins as inferred functional annotations, considering them as predicted GO terms for the hypothetical proteins. This network-based GO term prediction provided functional insights into these hypothetical proteins, further guiding their evaluation as potential vaccine candidates.

### 3.4. Vaccine Candidates

We have function predictions for all 83 hypothetical proteins from the three machine learning algorithms and the network analysis. After filtering the GO terms with the preset keywords and mapping back to the protein sequences, we got the vaccine candidate protein set from each algorithm. From the random forest prediction, we couldn’t get any candidates. But for AdaBoost, XGBoost and Network similarity predictions, we got respectively 24, 29 and 33 candidates. Notably, across all three candidate sets, we identified one common sequence with the ID “SeqID706,” suggesting that this protein may possess key functional characteristics making it more promising as a vaccine target.

Further analysis of “SeqID706” showed that it is linked to GO terms associated with both immune response activation and pathogen-host interaction, reinforcing its relevance. This convergence of predictions from multiple methods highlights “Se-qID706” as a strong candidate, warranting prioritized experimental validation.

## 4. Summary and Conclusion

This research investigates the functional prediction of hypothetical proteins in Aeromonas hydrophila, a pathogenic bacterium with significant implications for aquaculture. A. hydrophila presents a major challenge in fish farming due to its antibiotic resistance and pathogenicity, which notably affects warm, brackish environments. By identifying proteins associated with this pathogen’s virulence and survival mechanisms, this study seeks to contribute to vaccine development efforts that would mitigate its impact on aquaculture sustainability. Random Forest, XGBoost, and AdaBoost classifiers are used to predict hypothetical protein functions, complemented by network analysis. The research achieved functional predictions for all 83 hypothetical proteins in A. hydrophila using a combined ML and network analysis approach, which highlighted several candidate proteins for vaccine development. Notably, one protein, “SeqID706,” was consistently identified across models as a promising vaccine candidate due to its inferred role in immune response activation and host-pathogen interaction. The convergence of predictive results underscores the efficacy of ML and network analysis in uncovering potential immunogenic proteins, paving the way for targeted experimental validation.

